# Probability of Change in Life: amino acid changes in single nucleotide substitutions

**DOI:** 10.1101/729756

**Authors:** Kwok-Fong Chan, Stelios Koukouravas, Joshua Yi Yeo, Darius Wen-Shuo Koh, Samuel Ken-En Gan

## Abstract

Mutations underpin the processes in life, be it beneficial or detrimental. While mutations are assumed to be random in the bereft of selection pressures, the genetic code has underlying computable probabilities in amino acid phenotypic changes. With a wide range of implications including drug resistance, understanding amino acid changes is important. In this study, we calculated the probabilities of substitutions mutations in the genetic code leading to the 20 amino acids and stop codons. Our calculations reveal an enigmatic in-built self-preserving organization of the genetic code that averts disruptive changes at the physicochemical properties level. These changes include changes to start, aromatic, negative charged amino acids and stop codons. Our findings thus reveal a statistical mechanism governing the relationship between amino acids and the universal genetic code.

## INTRODUCTION

DNA is decoded in the frame of three bases at a time referred to as a codon (Crick et al., 1961). In eukaryotes, the host translational complex scans the mRNA for the start codon (i.e. AUG encoding for Methionine). Through the binding of amino acid bound-tRNA and codon pairing to anticodons on the tRNA, the codons are translated into amino acids based on the universal genetic code (Crick, 1968) with rare exceptions, and degeneracy at the third base, described as the Wobble Hypothesis (Crick, 1966). The majority of the common 20 amino acids are encoded by multiple codons (Crick et al., 1961; Lagerkvist, 1978) in the genetic code. Point mutations in DNA underpin many life processes and can occur as insertion, deletion and single nucleotide substitutions (SNS). Occurring at varying rates in life processes that range from hypermutations in the immune system e.g. antibody generation (Ling et al., 2018; Roth and Craig, 1998; Su et al., 2017) to disease development in cancer (Hanahan and Weinberg, 2011; Hollstein et al., 1991), and drug resistance (Chiang et al., 2018; Su et al., 2016; Weiss, 1993), mutations underpin change. Given that insertions and deletions cause frameshifts that are often detrimental in most cases, SNS are generally more frequent and are categorized into missense, nonsense and silent. Nonsense mutations often lead to disease states, for example, beta-thalassemia (Cao and Galanello, 2010) and cystic fibrosis (Tsui, 1992), amongst many others. However, non-conservative missense mutations can also cause diseases, as in the case of sickle cell anaemia (The International F.M.F. Consortium, 1997) where a missense mutation occurred on Glutamate (GAG to GTG) in the ß-globin gene.

Analysing the genetic code table, we found clear biases to the changes that mutations can achieve in limited mutation events. It is virtually impossible for Methionine (ATG) to mutate to a codon encoding for Proline (CCA, CCC, CCA, CCG) in a single SNS mutational event even though it is possible to become a Lysine (AAG) or Leucine (TTG). Such observations hint of an in-built probabilistic predisposition of innate mutational constraints. To address this, we analysed the probability of SNS induced changes in all 64 codons, with the aim of calculating the probable change outcomes of each codon.

## MATERIALS & METHODS

### Base probability of AA / stop signal in SNS mutations

The calculation of obtaining a specific codon during SNS mutations of T, A, C, G, respectively or when not specified is as follows in Equation 1:

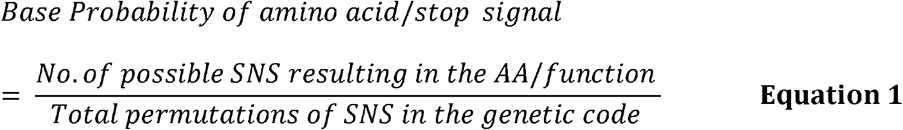

In the event where the resulting substituted base is known (the first four columns T, A, C and G of Figure 1), the following method is used to calculate the total permutations of SNS in the genetic code. The total number of possible SNS that can occur in the 64 codons is 144 (i.e. 3 codon positions × 64 codons = 192, and subsequently excluding 3 codon positions × 16 permutations for each pair of unmutated bases= 48, thus resulting in 144). The 16 permutations are excluded given mutations to self are not considered substitutions. Thus, if “A” was the resulting base on the first position of the codon “AXX” then the 16 other permutations are : {**A**AA, **A**AT, **A**AG, **A**AC, **A**TA, **A**TT, **A**TG, **A**TC, **A**GA, **A**GT, **A**GG, **A**GC, **A**CA, **A**CT, **A**CG, **A**CC}. In the case of Tyrosine (TAC or TAT), the number of possible T SNS encoding Tyrosine becomes 9 as follows: - first position - {**C**AT, **C**AC, **A**AT, **A**AC, **G**AT, **G**AC}; last position - {TA**C**, TA**A**, TA**G**} with a final probability of 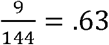.

**Figure 1.**
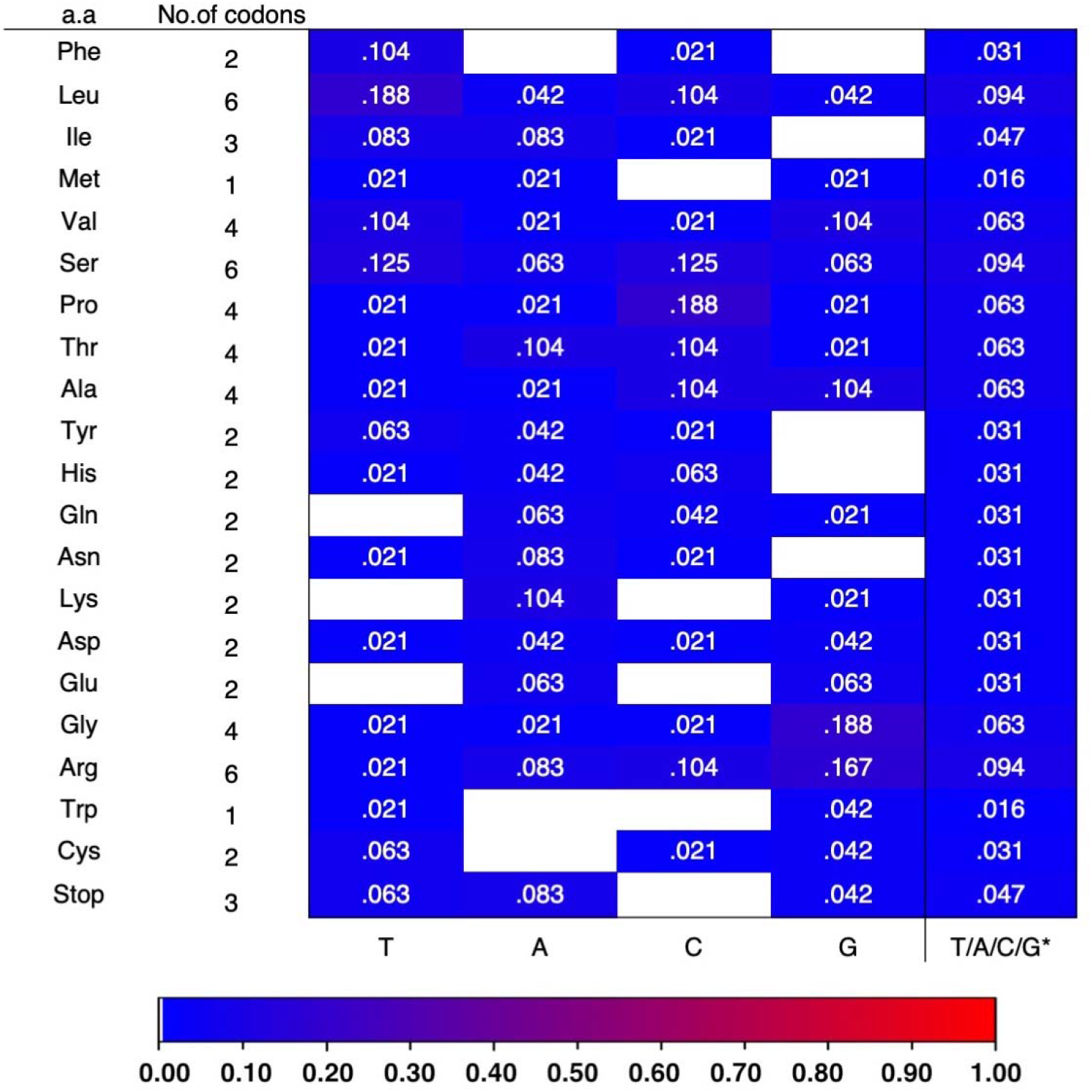
Base probability of AA and stop signal due to defined (T, A, C and G) / undefined (T/A/C/G) SNS mutations. Colour scheme: Blue to red based on increasing probability.

If the target base (T, A, G or C) mutation is unspecified (i.e. can be either of the four bases, rightmost column of Figure 1), the total combination becomes 64 codons × 3 codon positions × 3 bases = 576 possible permutations, excluding mutations to self. For Tyrosine, the number of possible SNS thereby encoding Tyrosine becomes 18 as follows: T - {**C**AT, **C**AC, **A**AT, **A**AC, **G**AT, **G**AC, TA**C**, TA**A**, TA**G**}; A - {T**T**T, T**T**C, T**C**T, T**C**C, T**G**T, T**G**C}; C - {TA**T**, TA**A**, TA**G**} with a final combined total probability of 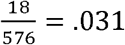.

### Probability of codon function change

In the probability of codon function change, the total number of possible changes of all codons for change is the number of codons encoding said amino acid/stop multiplied by 9 (reflecting the other 3 bases in the 3 positions of the codon) as shown in Equation 2.

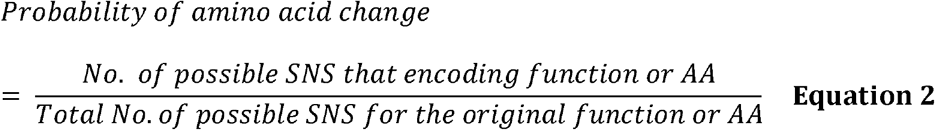

For example, for Arginine with 6 codons, there would be 6 × 9 = 54 possible changes in the total codons. Considering the mutations that can lead to Proline from Arginine codons as C**G**T, C**G**C, C**G**A, C**G**G, the probability of Arginine mutating to Proline would be 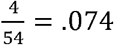

### Probability of Physicochemical property change

When amino acids are classified by their side chain hydrophobic and polar properties (Livingstone and Barton, 1993), the number of possible changes in all codons of the amino acids belonging to the specific physicochemical property group would be the denominator and calculated for specific groups using equation 3.

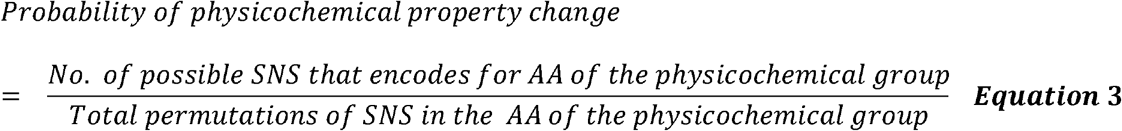

For example, the negatively charged amino acids, Aspartate and Glutamate are encoded by GAT, GAC and GAA, GAG, respectively. The number of possible SNS to 3 other bases in the 3 positions of the codon becomes 4 possible codons × 3 bases × 3 positions = 36 for the denominator. The codons that encode for negatively charged amino acid group when SNS occurs are 12 as follows - T: {GA**C**, GA**A**, GA**G**}, C : {GA**T**, GA**A**, GA**G**}, A : {GA**T**, GA**C**, GA**G**}, G : {GA**G**, GA**C**, GA**A**}, corresponding to 12 possible SNS. Thus, the probability of Aspartate and Glutamate mutating to other codons encoding Aspartate and Glutamate would be 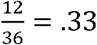.

## RESULTS

We found in-built predispositions in the genetic code when analysing the base probability of the 20 amino acids and stop codons occurring, underpinned by the four bases in specific locations of the codon (see Supplementary Figure S1 and S2A–C). This is illustrated where with a T in the first position of a codon, there is a bias towards codons encoding Serine (p = .25) over other amino acids (p < .2) as shown in Figure S1.

Studying the probability of a specific amino acid occurrence in the genetic code, predisposed changes to T, A, C or G lead to Leucine (p = .188), Lysine/Threonine (p = .104), Proline (p = .188) and Glycine (p =.188), respectively (Figure 1). This is due to the dominance of specific bases in the respective amino acid codons (Supplementary Table S3). Amino acids such as Serine, with a more balanced usage of the four bases, have more evened out probabilities compared with Leucine of also 6 codons. Alternatively, amino acid codons with a heavy bias towards specific bases, such as Proline (towards C) would have a higher probability of occurrence in a base C mutation event.

Calculating the probabilities of the 20 amino acids and stop codon to mutate to one another, there were interesting patterns observed when calculating the specific base changes (Supplementary Figure S4A-D). In the event of single G substitutions, Glycine will only mutate to itself. Such self-preservation probabilities are also observed for Phenylalanine in T substitutions, Lysine in A substitutions and Proline in C substitutions (Supplementary Figure S4A-D).

Probabilities of change to other amino acids were also not uniform (Supplementary Figure S4A–D and S5A–D). T mutations have a unique bias towards aromatic amino acids avoiding Glutamine, Lysine and Glutamate (Supplementary Figure S4A, Supplementary Figure S5A). Mutation to A, on the other hand, have higher probabilities to become stop codons and are less likely to lead to non-polar amino acids than other bases. Mutations to A also predisposes towards polar neutral or polar positive amino acids such as Lysine and Asparagine while avoiding Phenylalanine, Tryptophan and Cysteine (Supplementary Figure S4B, Supplementary Figure S5B). C mutations avert both start or stop codons, Lysine, Glutamate and Tryptophan, but has a strong bias for Proline (Supplementary Figure S4C, Supplementary Figure S5C), and G mutations have strongly pronounced biases towards Glycine and Arginine while avoiding Tyrosine, Isoleucine, Phenylalanine, Asparagine and Histidine (Supplementary Figure S4D, Supplementary Figure S5D).

When studying all 4 base SNS together, there is a pattern of self-preservation shown in the higher probabilities found diagonally in Figure 2 of no change. Self-bias was expectedly absent in amino acids encoded by 1 codon such as Methionine and Tryptophan.

**Figure 2.**
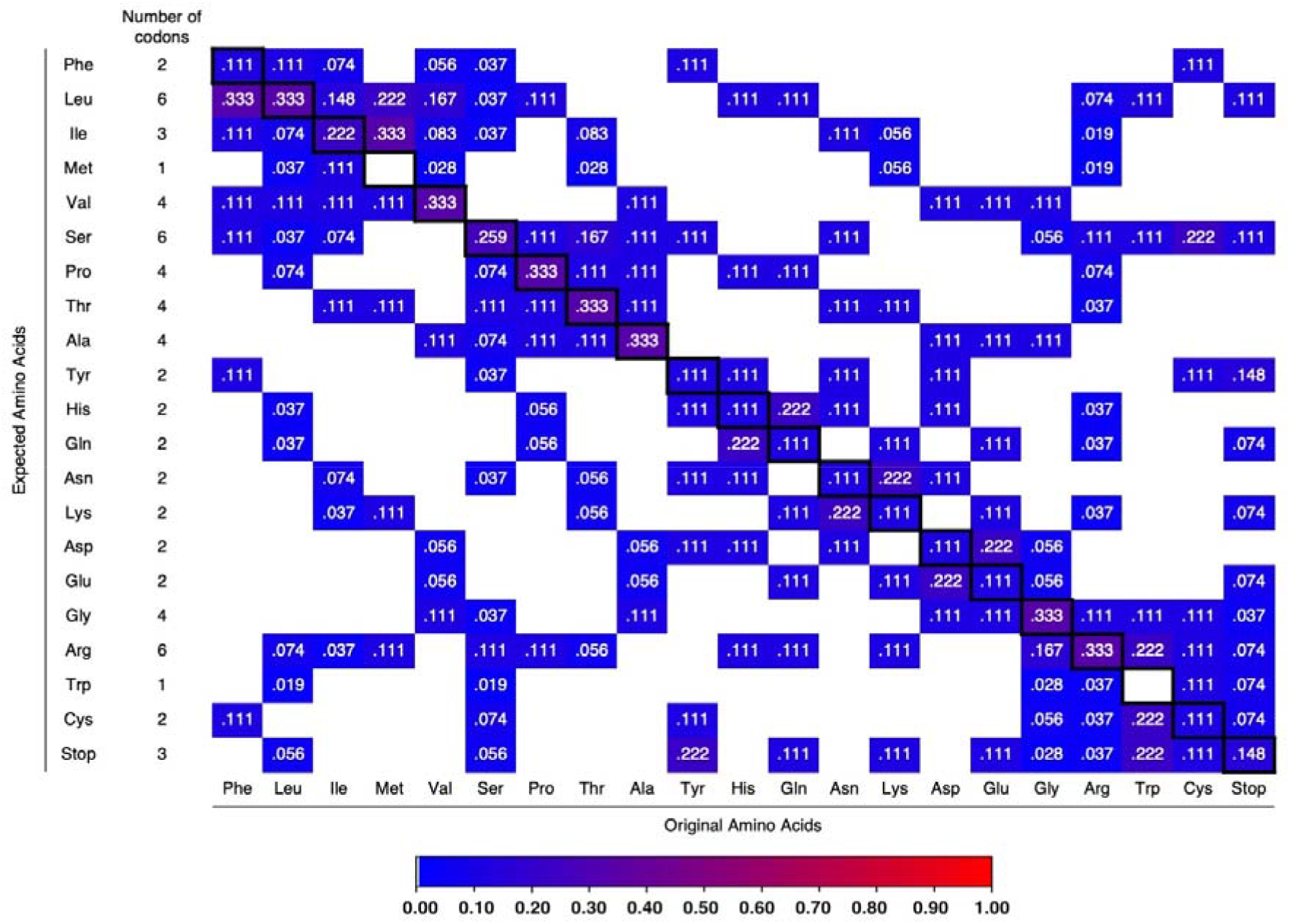
Probability of amino acids/stop signal changes. Colour scheme: Blue to red based on increasing probability. Probability tables of the amino acid change due to A, G, C, T mutations are separately calculated and shown in Supplementary Materials.

Although this relationship is not linear, the codon diversity plays a role in the probability of self-preservation; those amino acids with a more diverse range of codons have a decreased likelihood of retaining self, e.g. Arginine and Leucine with 6 codons, have the same self-preservation probability as the amino acids with 4 codons. On the other hand, Serine being more diverse despite also having 6 codons, had reduced self-preserving probability compared to Arginine and Leucine. Given that stop codons can tolerate changes in the second and third codon positions (TAA, TAG and TGA), the stop signal has lower probabilities of self-preservation compared to Isoleucine of also 3 codons with only changes in the third codon position (ATT, ATC, ATA) (Figure 2).

When grouped according to their physicochemical types, there are self-preserving biases towards silent and conservative changes within the same amino acid group (Figure 3). This self-bias applies to non-polar, polar and positive; the other groups have more levelled probabilities for non-conservative changes. Apart from stop codon self-preservation at p =.148 (Figure 3), the next highest pre-deposition towards stop codon (nonsense mutation) was by the aromatic group at p=.133. Specifically, Tryptophan and Tyrosine have the highest probabilities to a stop codon at p = .222 (Figure 2). Despite having many codons encoding them, non-polar amino acids have extremely low probabilities to change to a stop codon at p = .017 (Figure 3). Thus, intrinsic barriers against changes of amino acid and at the physicochemical levels are found intriguingly at the nucleic acid level.

**Figure 3.**
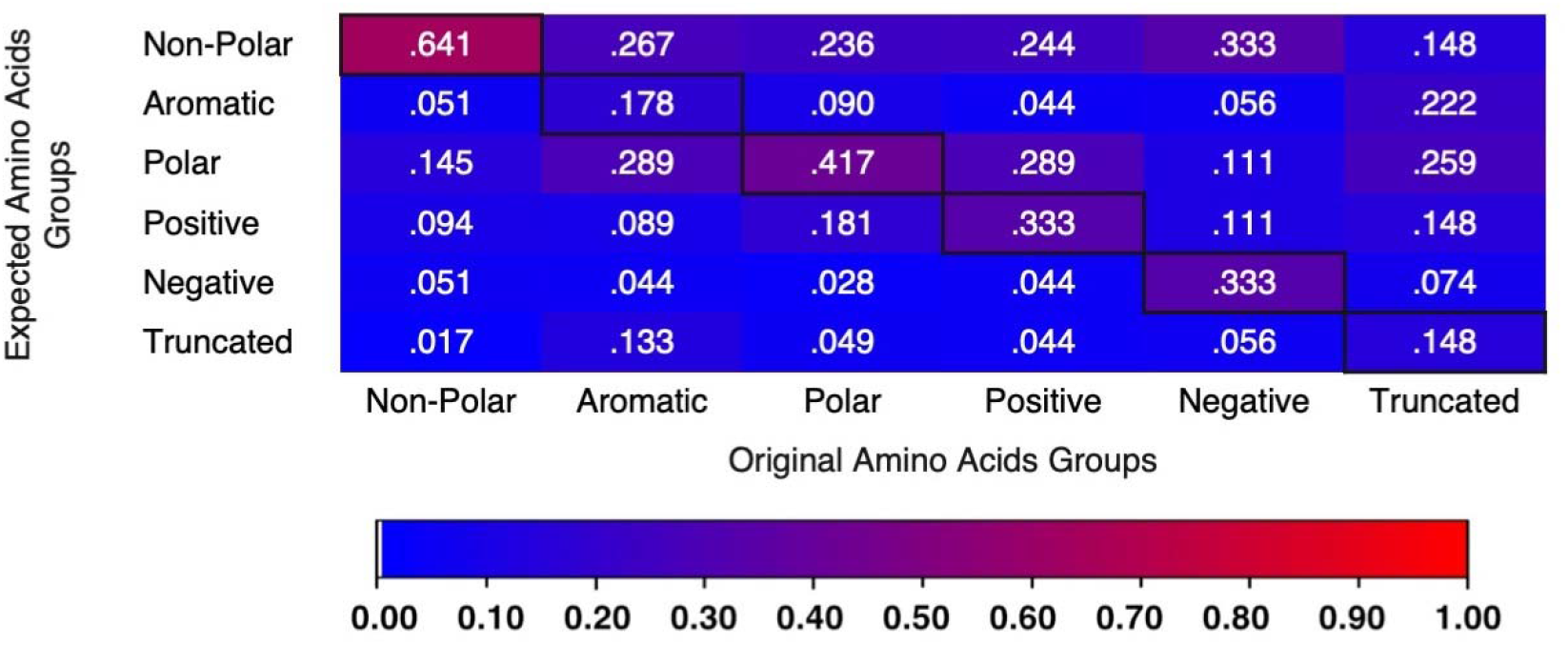
Probability of Physicochemical property changes (Livingstone and Barton, 1993). Colour scheme: Blue to red based on increasing probability.

## DISCUSSION

We found in-built intrinsic biases and barriers to drastic changes within the genetic code. Within single mutational events, there are fixed probabilities for the type of change in selection pressure-free conditions that are far from random.

In order to simulate the minimization of errors and ensure optimality (Błażej et al., 2018), the genetic code is shown to adopt a structure which relies on an internal and external system to avoid deleterious events (Figure 3) that could lead to the loss of genetic information. Instead, when driven towards other physicochemical group due to its internal probabilities, the genetic code allows fine-tuning between external biases of codon usage bias and mutation constraints, that mitigate errors during replicative processes (Zhu et al., 2003), protecting as much genetic information while being subjected to selection pressures.

While real-life applications cannot escape codon (Kurland, 1991) and mutational biases, they merely tilt the codon usage/mutation type of the organism rather than modify the innate “hard-coded” probabilities of specific codons mutating to those of amino acids. The predominance of specific tRNAs in species codon biases (Novoa et al., 2019) does not intrinsically change the probability of Glycine codons (GGG, GGA, GGT, GGC) to mutate to Methionine (ATG). Rather, the possible biases that could affect the intrinsic mutation probabilities would arise in the form of misincorporation of specific base pairs during replication/transcription. Factors for such misincorporation can be contributed by the varying availability of specific bases in the cell or biases elicited by enzymes e.g. Cysteine deaminases that lead to an increased misincorporation of U(s) in the presence of specific drugs (Goulian et al., 1980). Such biases, when analysed using the probability tables, allows an insight into the type of biological effects likely to be created, and even at the change of predisposition of further changes after the initial change.

As discussed in an article attempting to further optimize the genetic code using computer algorithms (Koonin and Novozhilov, 2009), such findings cannot reveal the origins of the code, or the selection steps to which the code could have arrived at the present robust, unique, and optimized state. In fact, the in-built self-preservation suggests that changes of the genetic code usage would be extremely rare in nature that is further confounded by the finite mutational events that can be incorporated into a population (Haldane, 1935) within a given time. With the constraint of the genetic code towards no change, preservation within the physicochemical group, or a slight trend towards non-polar amino acid codons (Figure 3), such predispositions at the nucleotide level supports the proposal of the rarity and optimality of the genetic code (Freeland and Hurst, 1998; Geyer and Madany Mamlouk, 2018; Wichmann and Ardern, 2019) in the universe, especially given the statistical self-bias. While certainly exciting, it would be unlikely given the odds, to find life in the universe that deviates greatly from the existing genetic code.

## FUTURE WORK

Since the genetic code has internal probabilities which extend beyond the scope of codon usage bias and mutational constraints, we foresee that these probabilities could be coupled with models which analyze the genetic code (Błażej et al., 2019), that would be useful for studying likely trajectories arising from known selection pressures in natural selection.

## CONCLUSION

We found statistical evidence that showed self-preservation and biases towards various amino acids in the genetic code table. In the event of substitution mutations, the highest probabilities are still conservative to steer away from aromatic, negative amino acids, and both start and stop codons. Such findings demonstrate self-preservation at the amino acid level occurring at the nucleotide level.

## ACKNOWLEDGEMENTS

This research was funded by the A*STAR core fund. We thank David Gunasegaran for his help in the calculations.

## AUTHOR STATEMENT

K.F.C. validated the manual calculation computationally. S.K. performed the calculations manually. S.K., K.F.C., J.Y.Y, W.S.D.K. and S.K.E.G. analysed the results and wrote the manuscript. S.K.E.G. conceived and supervised the study. All authors read and approved the final version of the manuscript.

## REFERENCES

Błażej, P., Fimmel, E., Gumbel, M., 2019. The Quality of Genetic Code Models in Terms of Their Robustness Against Point Mutations. Bulletin of Mathematical Biology 81, 2239–2257, doi:10.1007/s11538-019-00603-2.

Błażej, P., Wnętrzak, M., Mackiewicz, D., Mackiewicz, P., 2018. Optimization of the standard genetic code according to three codon positions using an evolutionary algorithm. PLOS ONE 13, e0201715, doi:10.1371/journal.pone.0201715.

Cao, A., Galanello, R., 2010. Beta-thalassemia. Genetics in Medicine 12, 61.

Chiang, R. Z.-H., Gan, S. K.-E., Su, C. T.-T., 2018. A computational study for rational HIV-1 non-nucleoside reverse transcriptase inhibitor selection and the discovery of novel allosteric pockets for inhibitor design. Bioscience reports 38, BSR20171113.

Crick, F. H., 1966. Codon—anticodon pairing: The wobble hypothesis. Journal of Molecular Biology 19, 548–555, doi:10.1016/S0022-2836(66)80022-0.

Crick, F. H., 1968. The origin of the genetic code. Journal of Molecular Biology 38, 367–379, doi:10.1016/0022-2836(68)90392-6.

Crick, F. H., Barnett, L., Brenner, S., Watts-Tobin, R. J., 1961. General nature of the Genetic Code for Proteins. Nature 192, 1227–1232, doi:10.1038/1921227a0.

Freeland, S. J., Hurst, L. D., 1998. The genetic code is one in a million. Journal of Molecular Evolution 47, 238–248.

Geyer, R., Madany Mamlouk, A., 2018. On the efficiency of the genetic code after frameshift mutations. PeerJ 6, e4825–e4825, doi:10.7717/peerj.4825.

Goulian, M., Bleile, B., Tseng, B., 1980. Methotrexate-induced misincorporation of uracil into DNA. Proceedings of the National Academy of Sciences 77, 1956–1960.

Haldane, J. B., 1935. The rate of spontaneous mutation of a human gene. Journal of Genetics 31, 317.

Hanahan, D., Weinberg, R. A., 2011. Hallmarks of cancer: the next generation. Cell 144, 646–674.

Hollstein, M., Sidransky, D., Vogelstein, B., Harris, C. C., 1991. p53 mutations in human cancers. Science 253, 49–53.

Koonin, E. V., Novozhilov, A. S., 2009. Origin and evolution of the genetic code: The universal enigma. IUBMB Life 61, 99–111, doi:10.1002/iub.146.

Kurland, C., 1991. Codon bias and gene expression. FEBS Letters 285, 165–169.

Lagerkvist, U., 1978. “ Two out of three”: an alternative method for codon reading. Proceedings of the National Academy of Sciences 75, 1759–1762.

Ling, W.-L., Lua, W.-H., Poh, J.-J., Yeo, J. Y., Lane, D. P., Gan, S. K.-E., 2018. Effect of VH–VL Families in Pertuzumab and Trastuzumab Recombinant Production, Her2 and FcγIIA Binding. Frontiers in immunology 9, 469.

Livingstone, C. D., Barton, G. J., 1993. Protein sequence alignments: a strategy for the hierarchical analysis of residue conservation. Bioinformatics 9, 745–756.

Novoa, E. M., Jungreis, I., Jaillon, O., Kellis, M., 2019. Elucidation of Codon Usage Signatures across the Domains of Life. Molecular Biology and Evolution 36, 2328–2339, doi:10.1093/molbev/msz124.

Roth, D. B., Craig, N. L., 1998. VDJ recombination: a transposase goes to work. Cell 94, 411–414.

Su, C. T.-T., Ling, W.-L., Lua, W.-H., Haw, Y.-X., Gan, S. K.-E., 2016. Structural analyses of 2015-updated drug-resistant mutations in HIV-1 protease: an implication of protease inhibitor cross-resistance. BMC Bioinformatics 17, 500.

Su, C. T.-T., Ling, W.-L., Lua, W.-H., Poh, J.-J., Gan, S. K.-E., 2017. The role of antibody Vκ framework 3 region towards antigen binding: effects on recombinant production and protein L binding. Scientific reports 7, 3766.

The International F.M.F. Consortium, 1997. Ancient Missense Mutations in a New Member of the RoRet Gene Family Are Likely to Cause Familial Mediterranean Fever. Cell 90, 797–807, doi:10.1016/S0092-8674(00)80539-5.

Tsui, L.-C., 1992. The spectrum of cystic fibrosis mutations. Trends in Genetics 8, 392–398.

Weiss, R. A., 1993. How does HIV cause AIDS? Science 260, 1273–1279.

Wichmann, S., Ardern, Z., 2019. Optimality in the standard genetic code is robust with respect to comparison code sets. Biosystems 185, 104023, doi:10.1016/j.biosystems.2019.104023.

Zhu, C.-T., Zeng, X.-B., Huang, W.-D., 2003. Codon Usage Decreases the Error Minimization Within the Genetic Code. Journal of Molecular Evolution 57, 533–537, doi:10.1007/s00239-003-2505-7.

