## Supplementary for "Probability of Change in Life: amino acid changes in single nucleotide substitutions"

**SUPPLEMENTARY INFORMATION**

**Probability of amino acid occurrence in the genetic code**

With specific bases (T, A, C, G) at defined and undefined codon positions, the probabilities of the encoding amino acid is calculated using Supplementary Equation 1, where the total permutations with specific bases occurring at defined and undefined codon positions would be the denominator.

$$Probability ofamino acid given specific base = \frac{No. of codons with specific base at codon position}{Total permutations for specific base at vaious codon position} \mathbf{Supp. Equation 1}$$

For example, the number of codons with T at the codon 1st position leading to Phenylalanine is 2 (**T**TT, **T**TC). There are also 16 possible permutations within the genetic code with T in the 1st position (Top row of codon table). Thus, the calculated probability of Phe given that T is located at 1st position in the codon is $\frac{2}{16}=0.125$.

For an undefined location of T in the codon, the total possible permutations is 16 x 3 positions = 48 in the denominator (1st – Top row, 2nd –most left column and 3rd – first codon of every 4 rows in codon table). The possible permutations with T leading to Phe are at the 1st position (**T**TT, **T**TC), 2nd position (T**T**T, T**T**C) and the 3rd position (TT**T**), which corresponds to the numerator. Thus, the probability of T in undefined position leading to Phenylalanine is $\frac{5}{48}=0.104$.

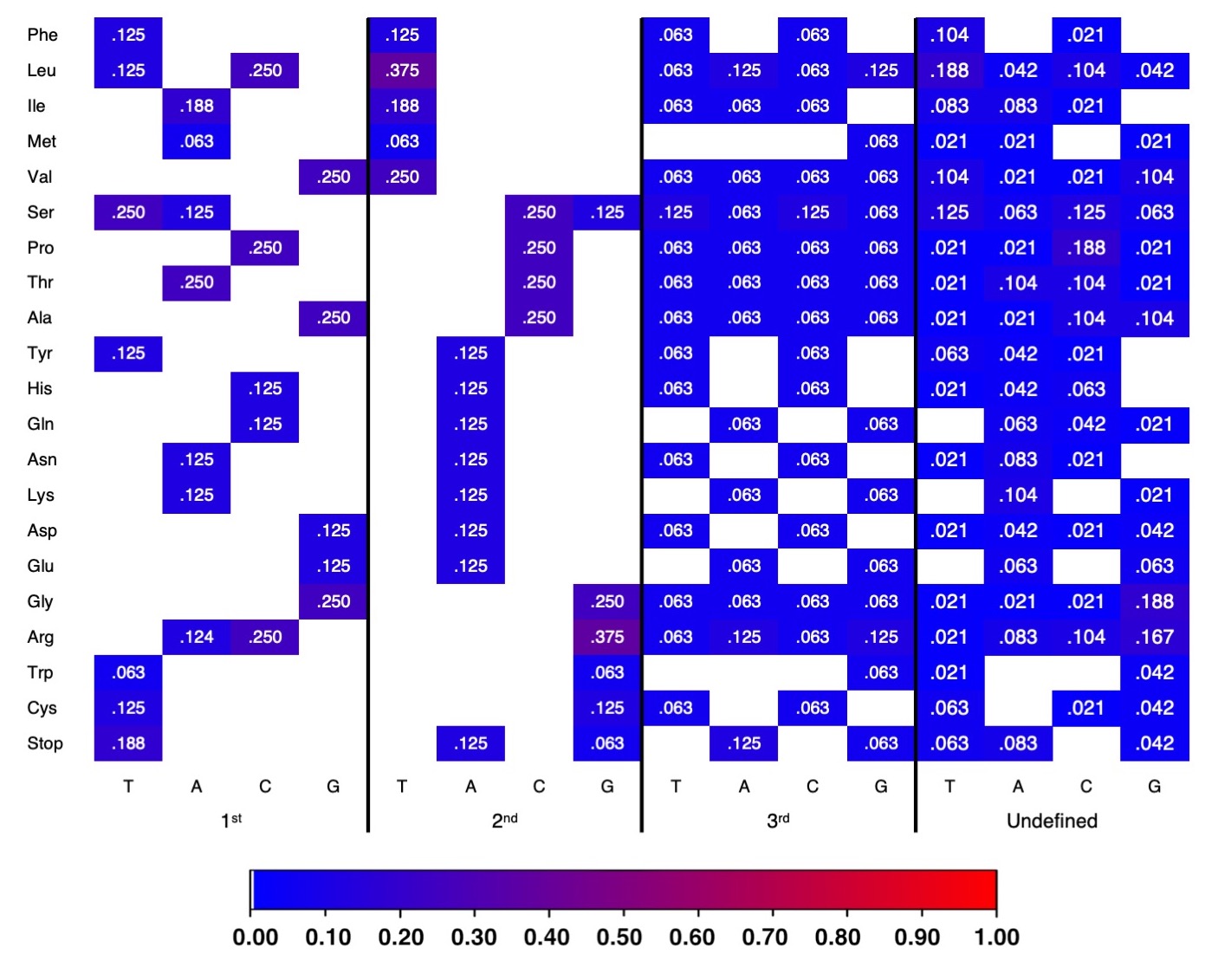

**Supplementary Figure S1.** Probability of the amino acid occurrence in the genetic code. Further calculations are in Supplementary Figure S2A-C. Colour scheme: Blue to red based on increasing probability with white being p= 0.

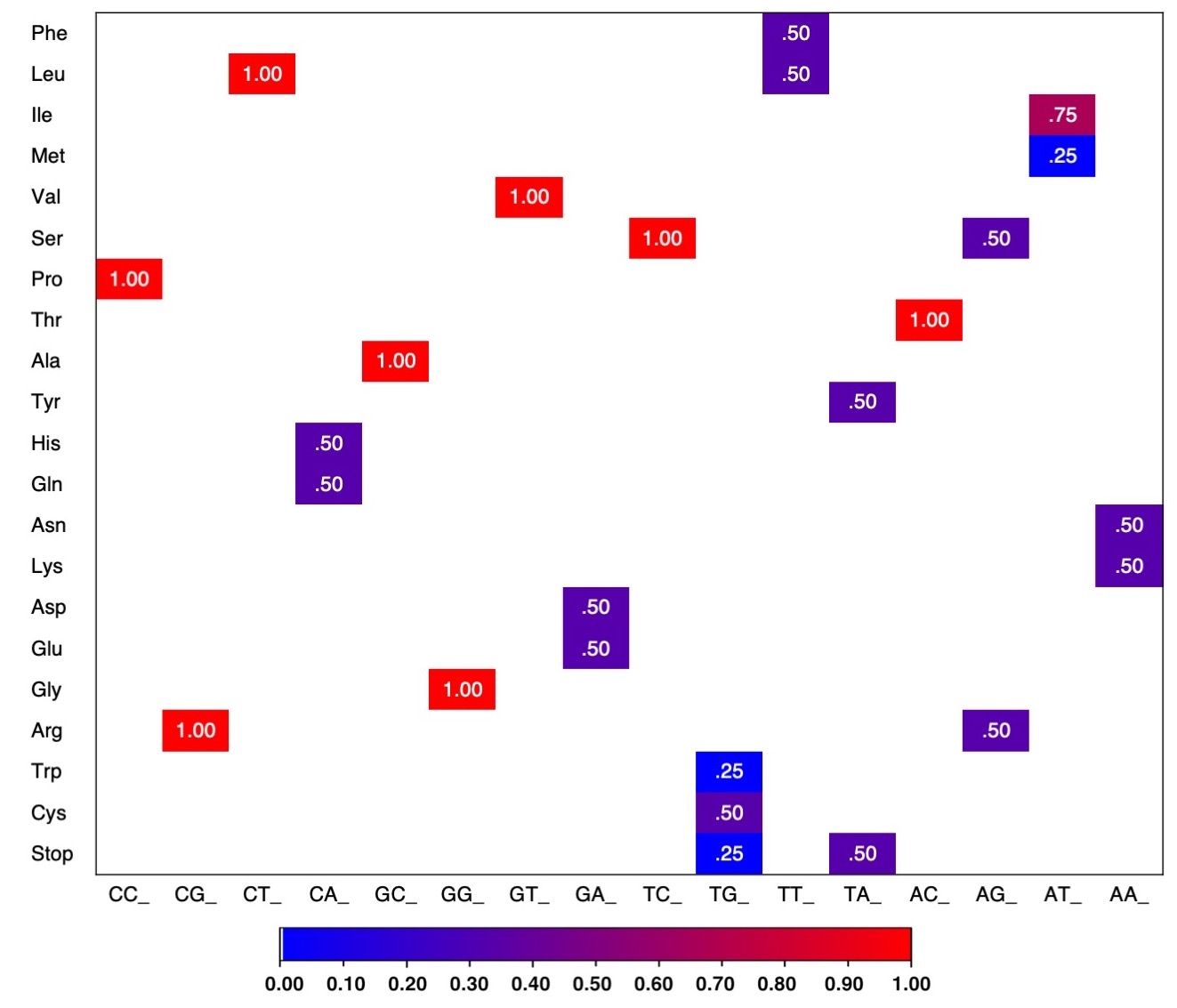

**Supplementary Figure S2A**. Probability of the specific amino acid and stop codon occurrences when the first two codon nucleotides are known.

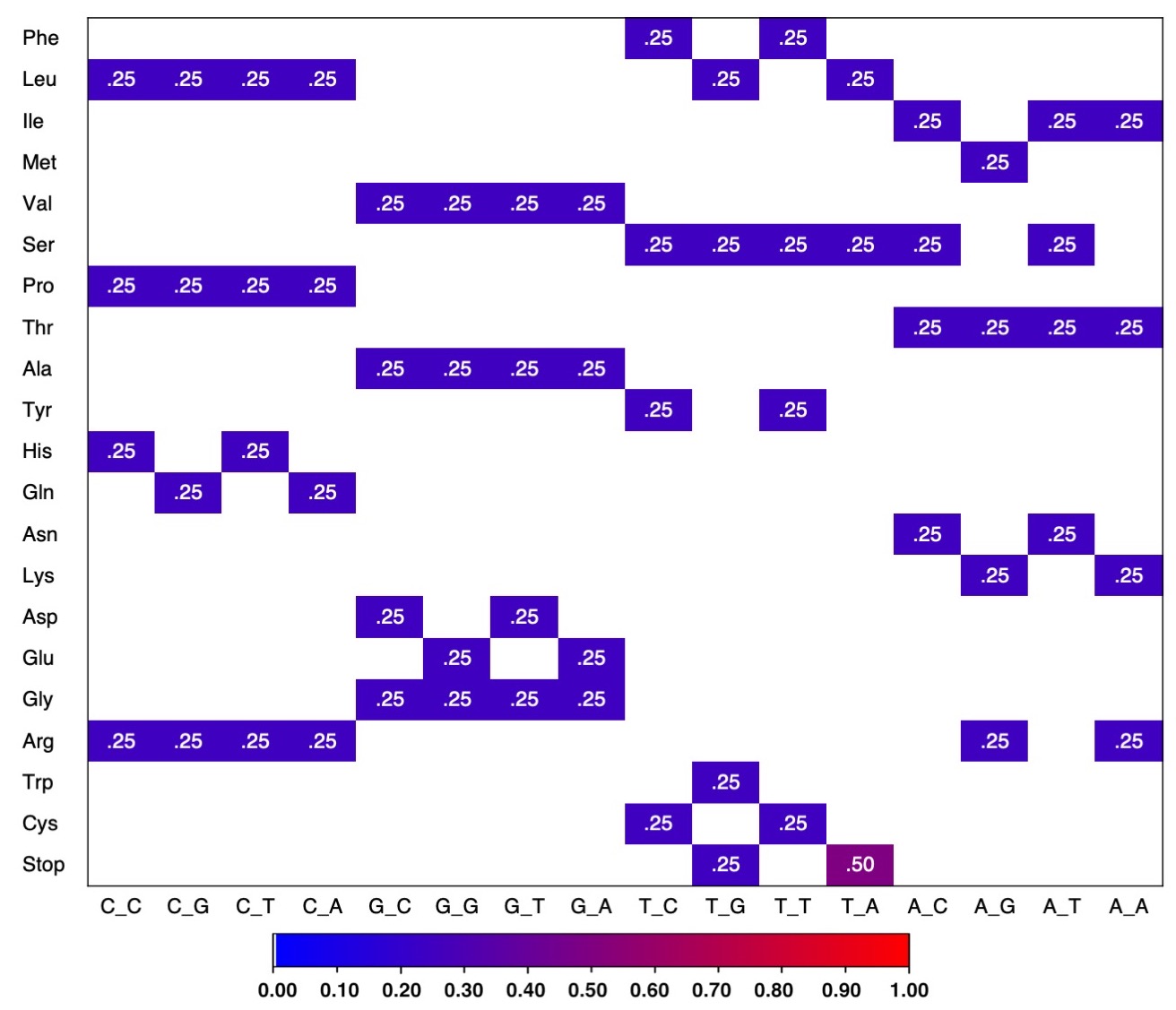

**Supplementary Figure S2B.** Probability of the specific amino acid and stop codon occurrences in the respective known 1st and 3rd codon nucleotides.

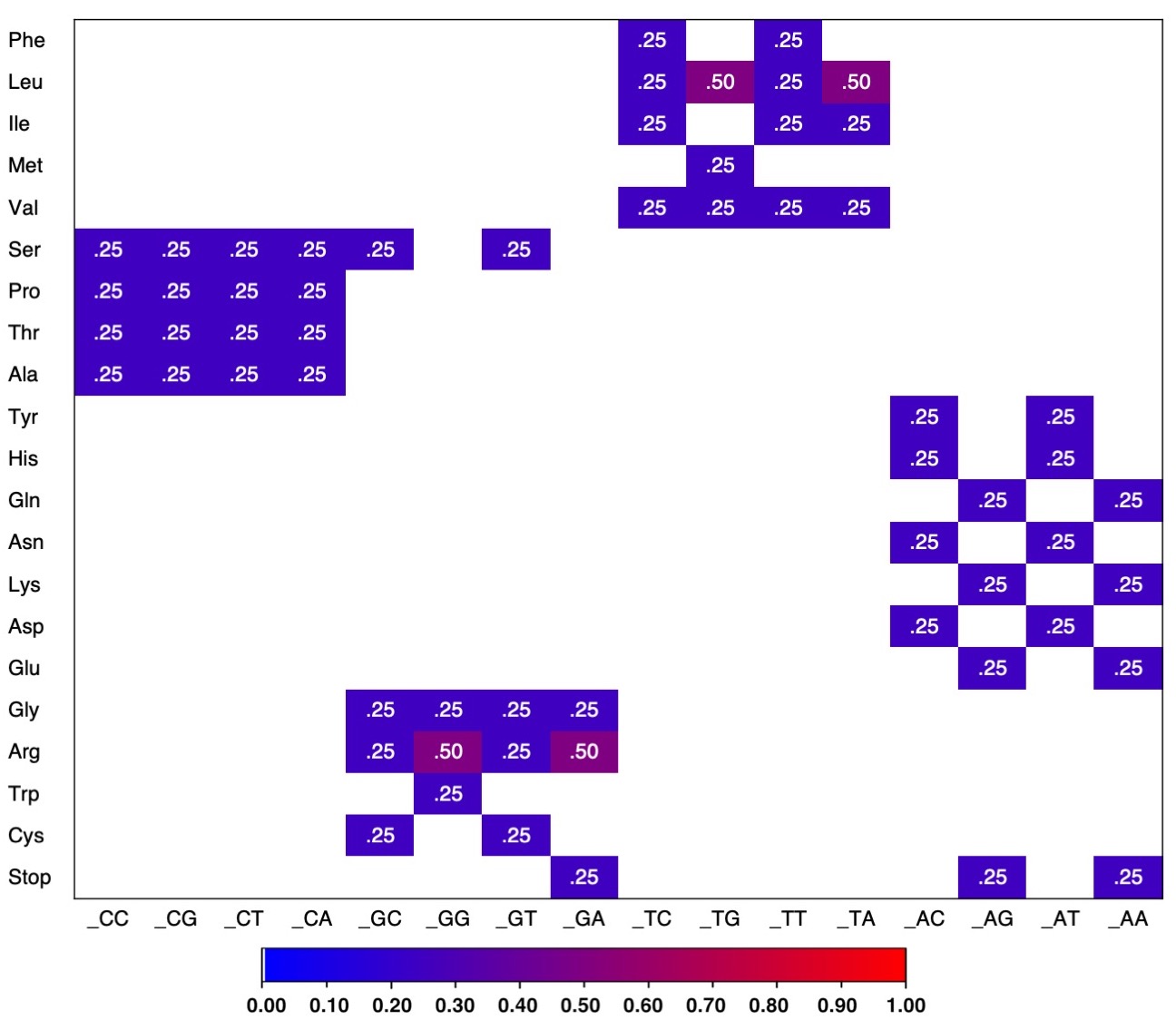

**Supplementary Figure S2C.** Probability of the specific amino acid and stop codon occurrences in the respective known 2nd and 3rd codon nucleotides.

**Occurrences of amino acid in specific base and codon positions**

The total number of possible permutations in undefined codon positions for a specific base is given by the number of occurrences with the specific base in 1st, 2nd, and 3rd positions.

For example, there are 2 occurrences with A in the 1st position (**A**AT, **A**AC), 2 occurrences of A in the 2nd position (A**A**T, A**A**C) and 1 occurrence of T and C in the 3rd position respectively (AA**T**, AA**C**).

**Supplementary Table S3.** Occurrences of the respective amino acid (a.a) in conditions of known bases at the various codon positions.

|  | 1^st^ | | | | 2^nd^ | | | | 3^rd^ | | | | Undefined* | | | |
| --- | --- | --- | --- | --- | --- | --- | --- | --- | --- | --- | --- | --- | --- | --- | --- | --- |
| **a.a** | T | A | C | G | T | A | C | G | T | A | C | G | T | A | C | G |
| **Phe** | 2 |  |  |  | 2 |  |  |  | 1 |  | 1 |  | 5 |  | 1 |  |
| **Leu** | 2 |  | 4 |  | 6 |  |  |  | 1 | 2 | 1 | 2 | 9 | 2 | 5 | 2 |
| **Ile** |  | 3 |  |  | 3 |  |  |  | 1 | 1 | 1 |  | 4 | 4 | 1 |  |
| **Met** |  | 1 |  |  | 1 |  |  |  |  |  |  | 1 | 1 | 1 |  | 1 |
| **Val** |  |  |  | 4 | 4 |  |  |  | 1 | 1 | 1 | 1 | 5 | 1 | 1 | 5 |
| **Ser** | 4 | 2 |  |  |  |  | 4 | 2 | 2 | 1 | 2 | 1 | 6 | 3 | 6 | 3 |
| **Pro** |  |  | 4 |  |  |  | 4 |  | 1 | 1 | 1 | 1 | 1 | 1 | 9 | 1 |
| **Thr** |  | 4 |  |  |  |  | 4 |  | 1 | 1 | 1 | 1 | 1 | 5 | 5 | 1 |
| **Ala** |  |  |  | 4 |  |  | 4 |  | 1 | 1 | 1 | 1 | 1 | 1 | 5 | 5 |
| **Tyr** | 2 |  |  |  |  | 2 |  |  | 1 |  | 1 |  | 3 | 2 | 1 |  |
| **His** |  |  | 2 |  |  | 2 |  |  | 1 |  | 1 |  | 1 | 2 | 3 |  |
| **Gln** |  |  | 2 |  |  | 2 |  |  |  | 1 |  | 1 |  | 3 | 2 | 1 |
| **Asn** |  | 2 |  |  |  | 2 |  |  | 1 |  | 1 |  | 1 | 4 | 1 |  |
| **Lys** |  | 2 |  |  |  | 2 |  |  |  | 1 |  | 1 |  | 5 |  | 1 |
| **Asp** |  |  |  | 2 |  | 2 |  |  | 1 |  | 1 |  | 1 | 2 | 1 | 2 |
| **Glu** |  |  |  | 2 |  | 2 |  |  |  | 1 |  | 1 |  | 3 |  | 3 |
| **Gly** |  |  |  | 4 |  |  |  | 4 | 1 | 1 | 1 | 1 | 1 | 1 | 1 | 9 |
| **Arg** |  | 2 | 4 |  |  |  |  | 6 | 1 | 2 | 1 | 2 | 1 | 4 | 5 | 8 |
| **Trp** | 1 |  |  |  |  |  |  | 1 |  |  |  | 1 | 1 |  |  | 2 |
| **Cys** | 2 |  |  |  |  |  |  | 2 | 1 |  | 1 |  | 3 |  | 1 | 2 |
| **Stop** | 3 |  |  |  |  | 2 |  | 1 |  | 2 |  | 1 | 3 | 4 |  | 2 |

**Probability table of amino acid (a.a.) changes with a base (T, A, C, G) substitution.**

The method for calculating the probabilities can be found in the main manuscript Materials and Methods

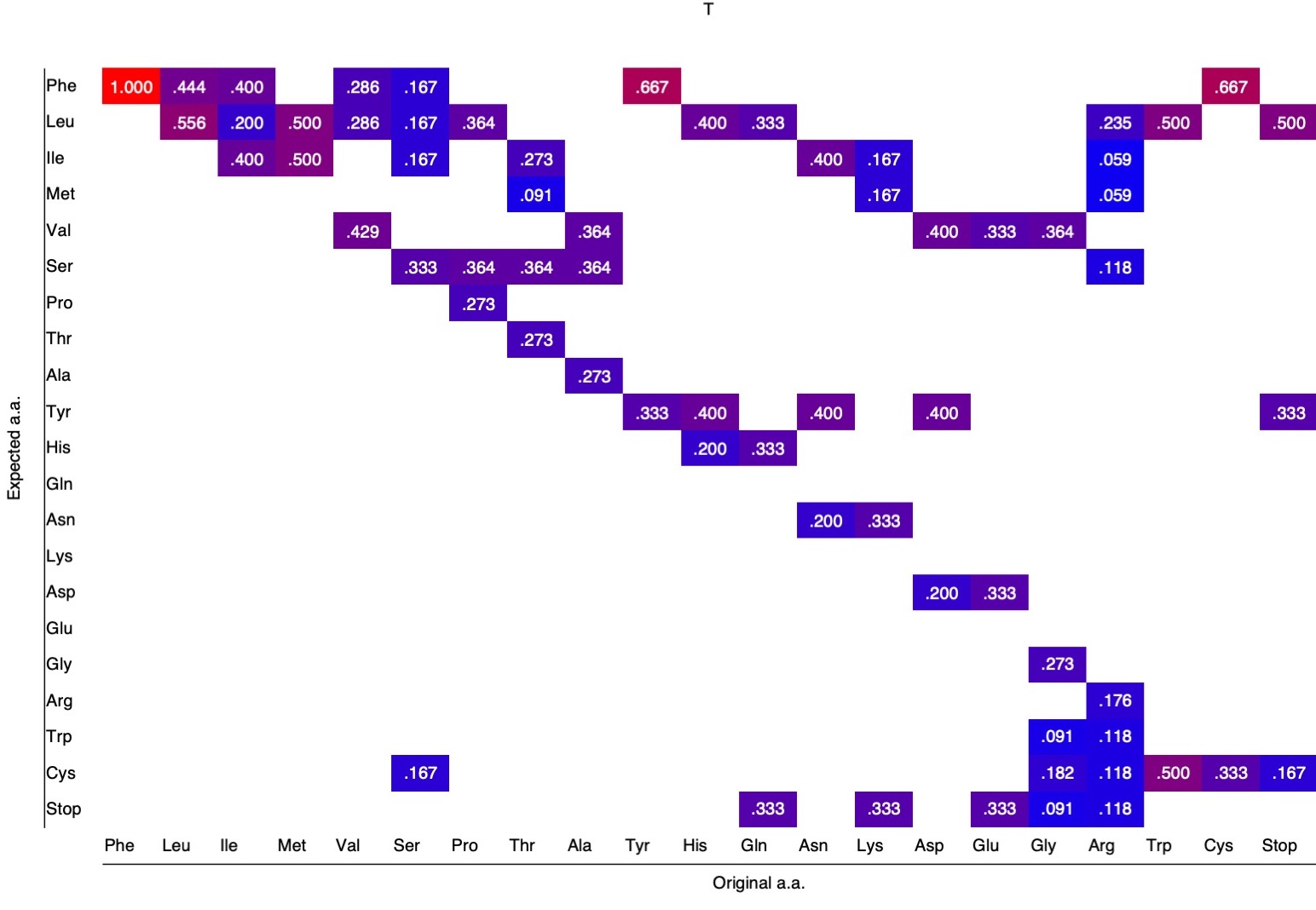

**Supplementary Figure S4A.** Probability table of amino acid (a.a.) changes with a T substitution.

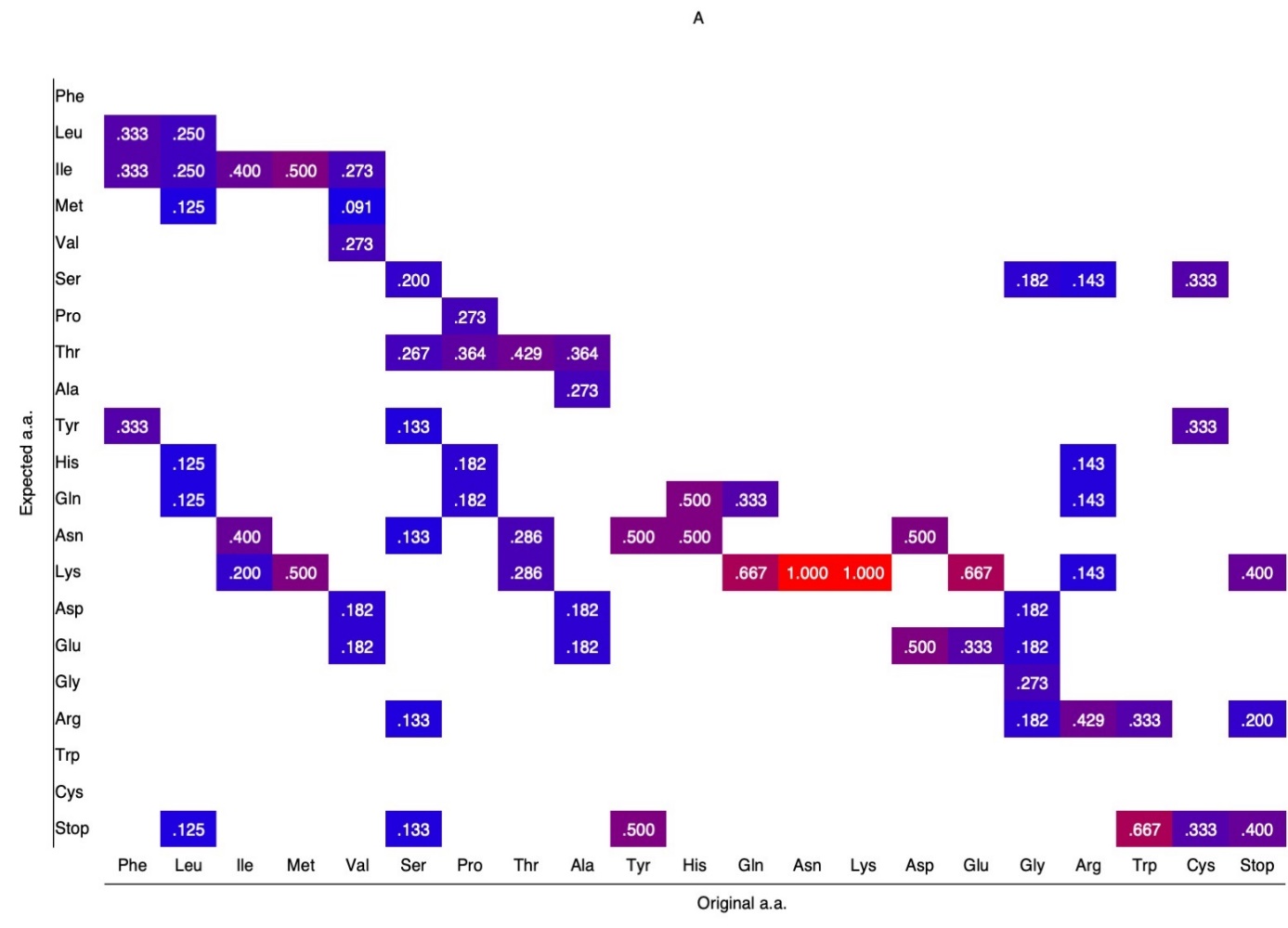

**Supplementary Figure S4B.** Probability table of amino acid (a.a.) changes with an A substitution.

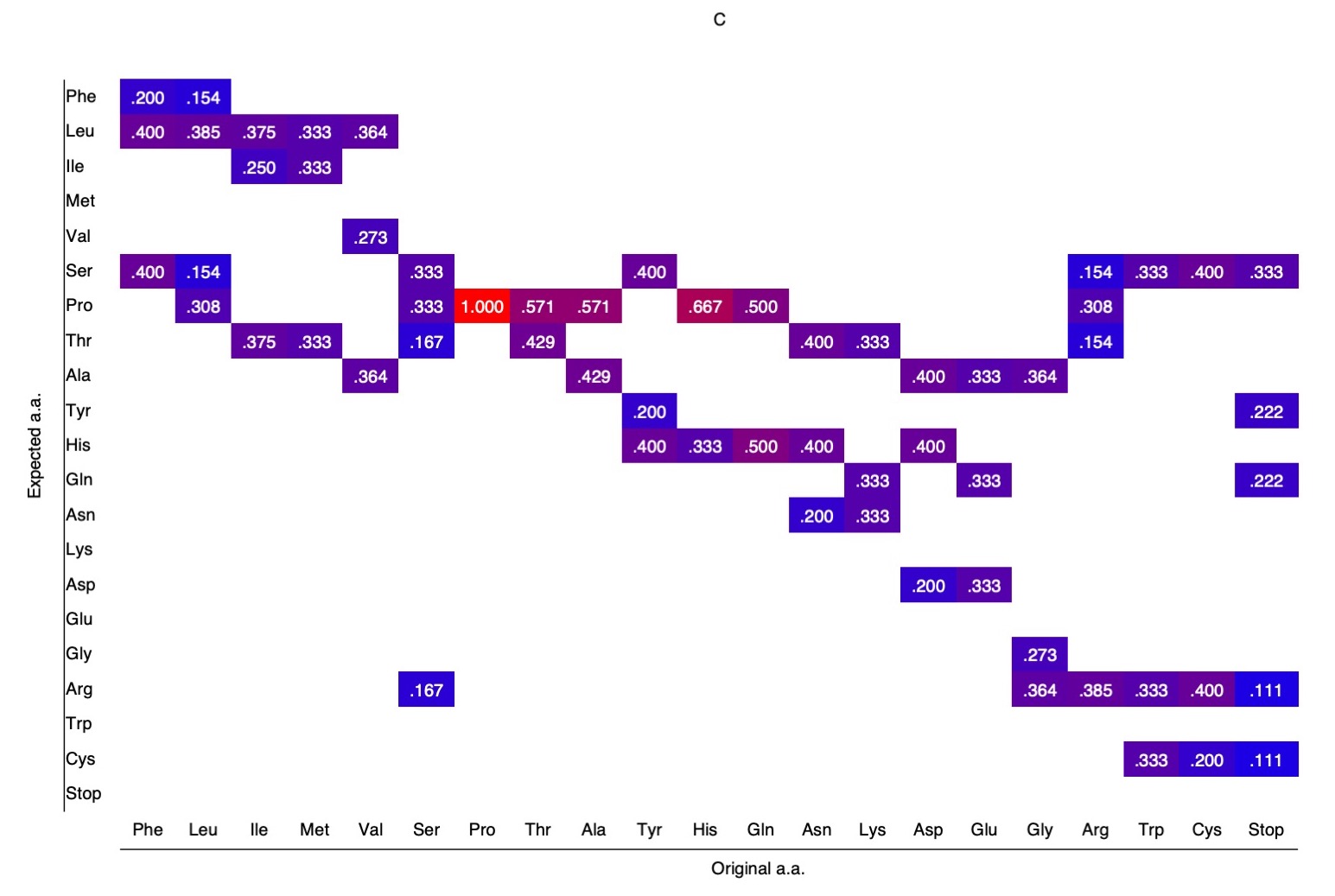

**Supplementary Figure S4C.** Probability table of amino acid (a.a.) changes with a C substitution.

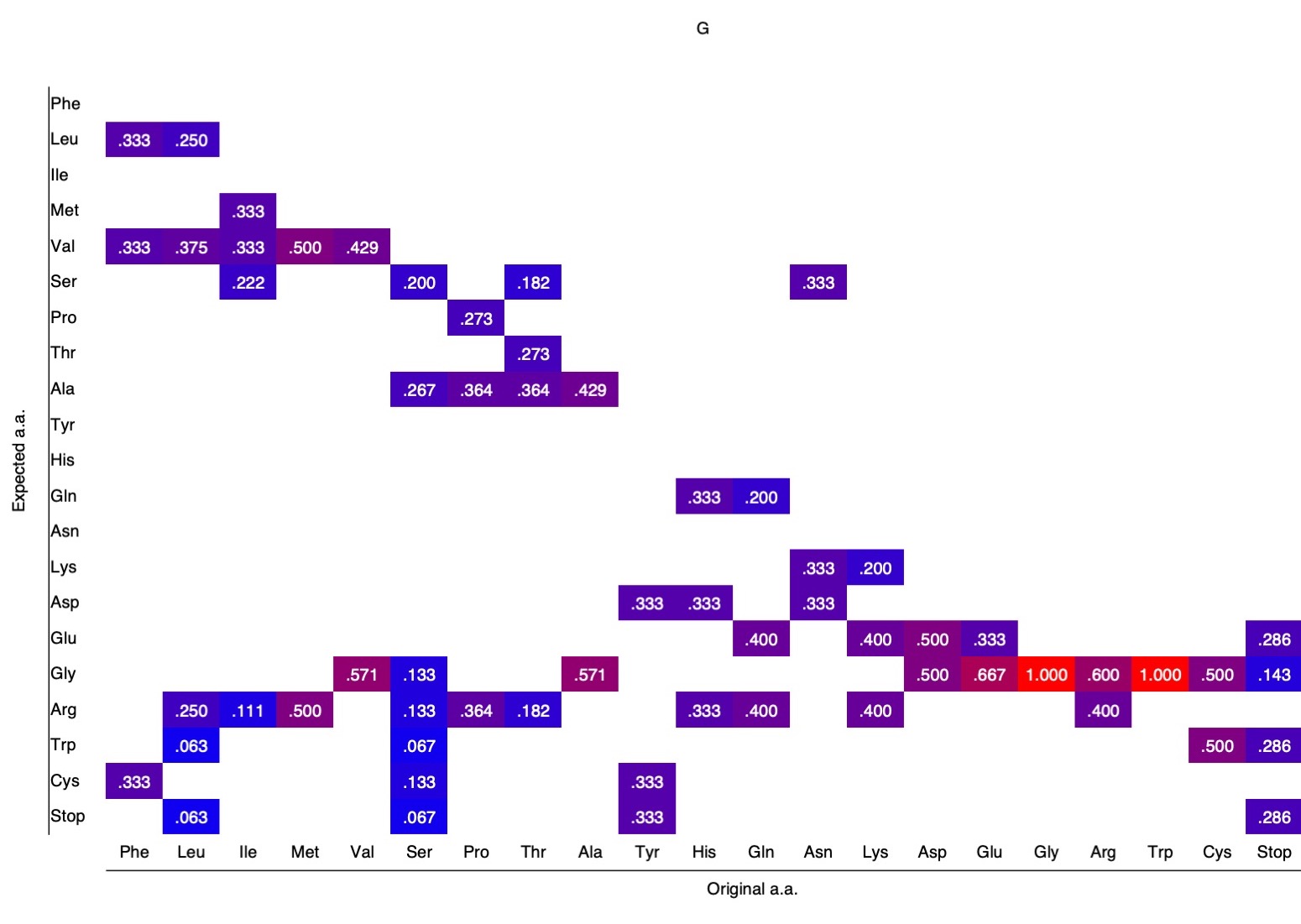

**Supplementary Figure S4D.** Probability table of amino acid (a.a.) changes with a G substitution.

**Probability of physicochemical type change with a base (T, A, C, G) substitution.**

The method for calculating the probabilities can be found in the main manuscript Materials and Methods under “Probability of Physicochemical property changes”

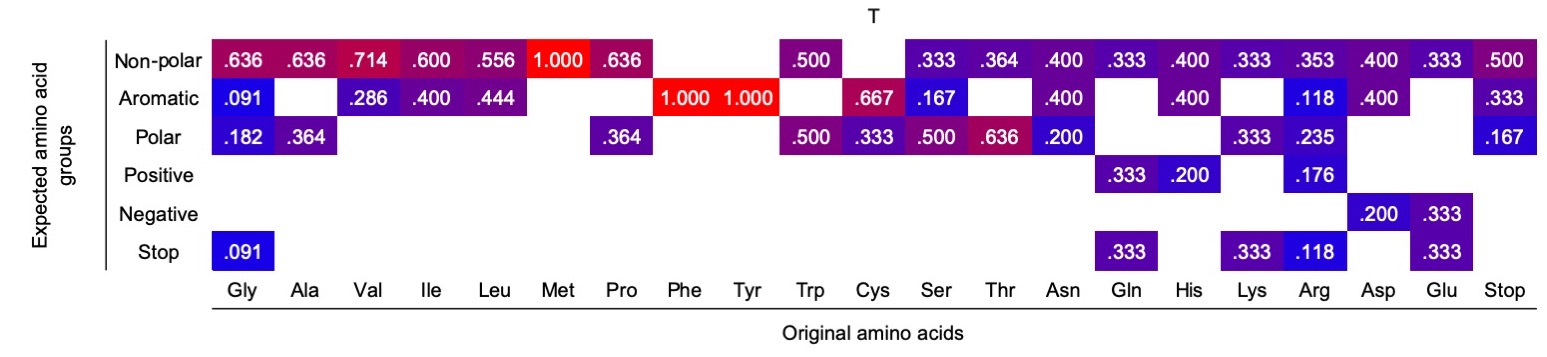

**Supplementary Figure S5A.** Probability of physicochemical type change with a T substitution.

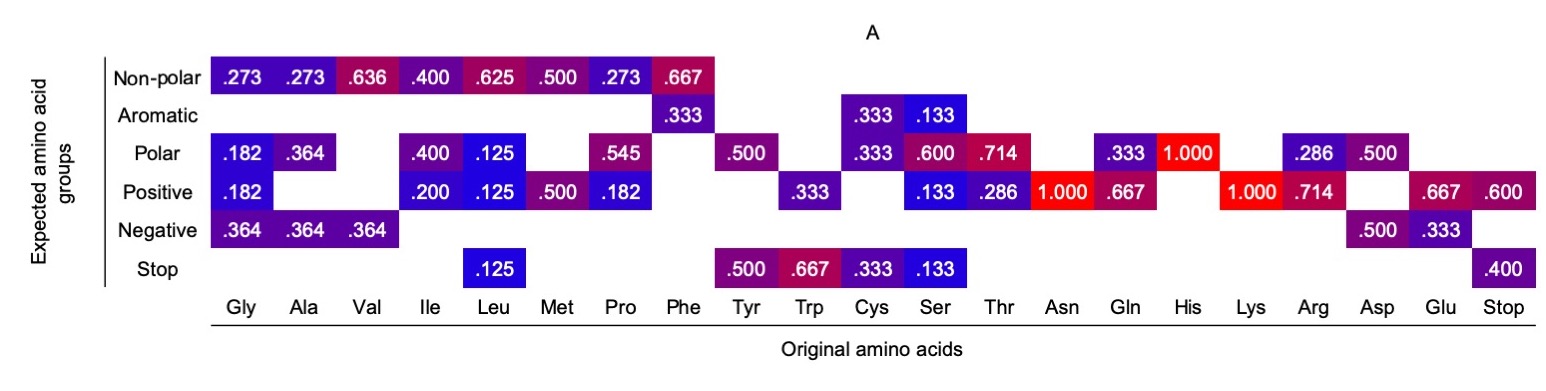

**Supplementary Figure S5B.** Probability of physicochemical type change with an A substitution.

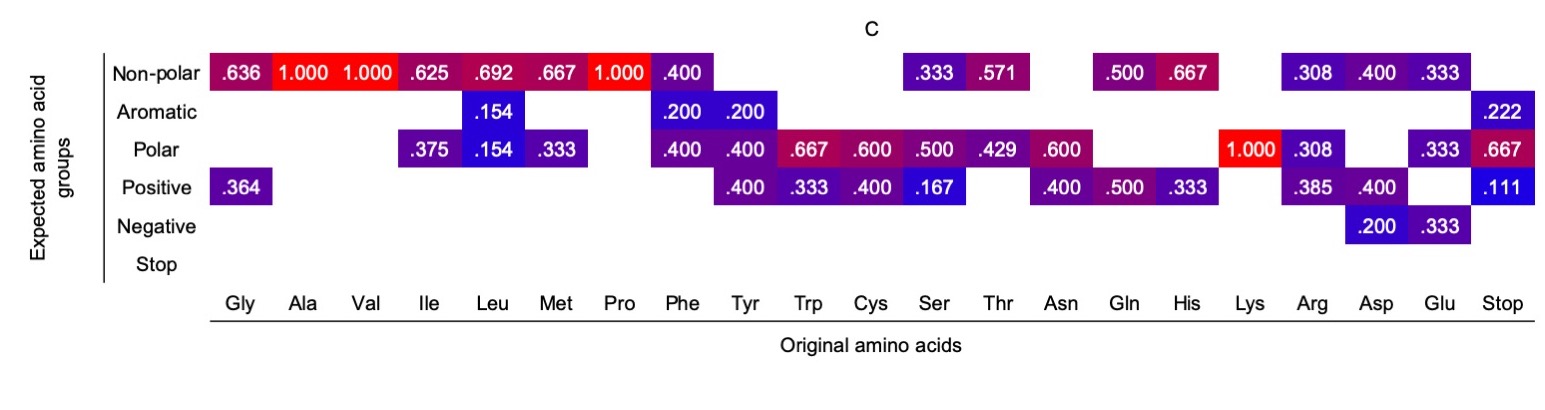

**Supplementary Figure S5C.** Probability of physicochemical type change with a C substitution.

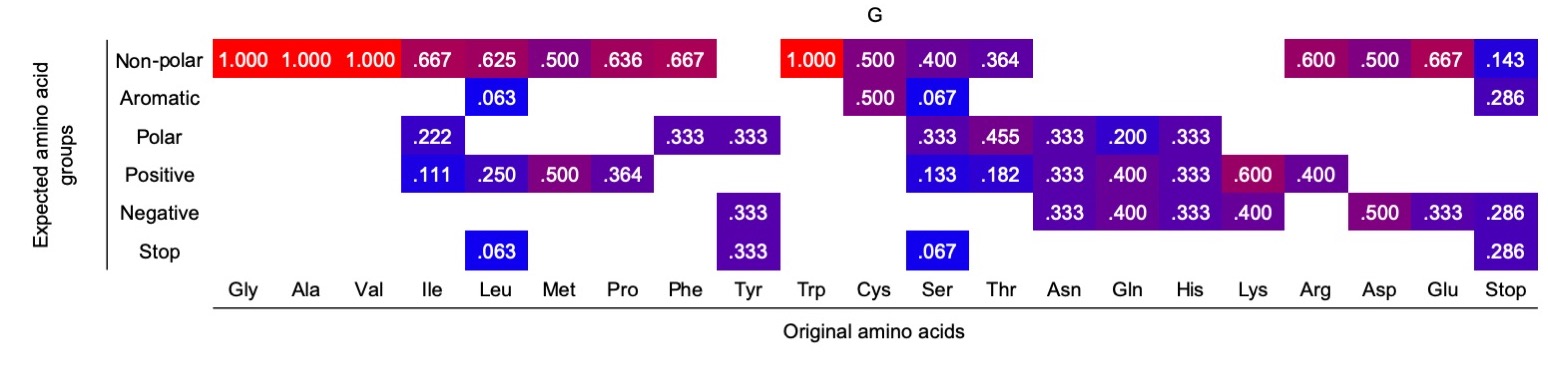

**Supplementary Figure S5D.** Probability of physicochemical type change with a G substitution
